# Review of histopathologic findings of endometrial biopsy done at a tertiary referral hospital in urban Ethiopia

**DOI:** 10.1101/2020.04.16.044610

**Authors:** Lemi B Tolu, Solomon Kabtamu, Mustefa Abedella, Garumma Tolu Feyissa, Elias Senbeto

## Abstract

**Background:** An endometrial sampling procedure is a gold standard for the diagnostic evaluation of women with abnormal uterine bleeding (AUB) with primary aim to identify the cause, especially to determine whether carcinoma or premalignant lesions are present. The objective of the study is therefore to review endometrial biopsy specimen histopathological findings and associated factors at a tertiary referral hospital in urban Ethiopia.

**Methodology:** This is a two years retrospective analysis of cases of patients for whom endometrial biopsy was taken at a territory referral hospital in urban Ethiopia. Odd’s ratios, 95% confidence interval and p-value set at 0.05 were used to determine the statistical significance of the associations.

**Results:** A total of 277 patient records were analyzed and mean and the median age of the study patients were 41.89 and 40.00 years respectively. The commonest histopathologic finding was endometrial polyp 66 (23.83%), followed by proliferative endometrium 47 (16.97%) and secretory endometrium 25(9.03%). Endometrial hyperplasia, endometrial malignancy, and pregnancy complications were reported in 9 (3.25%), 13 (4.69%), and 25 (9.03%) of cases respectively. Endometritis was detected in 20(7.22%), while Tuberculous (TB) endometritis was reported in 3(1.08%) of cases only. Inconclusive and inadequate sample was reported in 30 (10.83%) and 34 (12.27%) of cases respectively. Endometrial polyp was associated with 40-49 years of age (OR= 12.56, 95% CI: 2.58-61.23).

**Conclusions:** Endometrial polyp was the commonest histopathologic finding followed by proliferative and secretory endometrium respectively. Rate of sample inadequacy is higher than most of the study reports which mandates to improve sampling technique to increase sample adequacy and routine transvaginal ultrasound for endometrial evaluation especially for those postmenopausal women to decide the necessity of endometrial sampling.

## Background

An endometrial biopsy is the removal of a small piece of tissue from the endometrium, which is the lining of the uterus (1-3). This tissue sample can show cell changes due to abnormal tissues or variations in hormone levels. The preferred method is an outpatient office-based endometrial biopsy, usually without anesthesia or cervical dilation. For the office procedure, a 2–4 mm diameter plastic cannula, with a plunger within, is used and is reliable and is well accepted by the patient (1-7).

Sharp curettage (D& C) alone or in combination with hysteroscope might also be required in case simple office sampling techniques failed to provide sufficient diagnostic information or if abnormal bleeding persists after normal histopathologic findings (2, 3).

An endometrial sampling procedure is a gold standard for the diagnostic evaluation of women with abnormal uterine bleeding (AUB) with primary aim to identify the cause, especially to determine whether carcinoma or premalignant lesions are present(3). The incidence and risk of endometrial cancer increase with age and AUB is noted in 80 to 90 percent of women with endometrial cancer. Thus, in postmenopausal women, the need to exclude cancer intensifies, and endometrial biopsy is typically indicated. Of premenopausal women with endometrial neoplasia, most are obese or have chronic anovulation or both. Thus, women with AUB in these two groups also warrant exclusion of endometrial cancer(8-11).

Mapping and documenting the result of endometrial sampling is very important to associate patient clinical symptoms and histopathologic finding. This is important to stratify the management of patients based on documented evidences. Accordingly, retrospective review of endometrial pathology in Romania among patients with postmenopausal bleeding reported atrophy of the endometrium to be the predominant finding (41.85%). Endometrial polyps, endometrial adenocarcinoma and endometrial hyperplasia were reported in about 12.02%, 27.52%, and 15.50% of women respectively(11). Similarly another four years retrospective review of endometrial pathology in the University Hospital Bucharest endometrial hyperplasia’s without atypia represented nearly half of the pathologies (12).

As far as we know there are very few studies reported in our country Ethiopia from Tikur Anbessa Specialized Teaching Hospital (TASTH) before seven years. The first was retrospective review on 448 patient records and reported that the commonest pattern in these patients was normal cycling endometrium (44.9%) (13). The second study was conducted to determine the prevalence of endometrial tuberculosis among patients who undergoes endometrial sampling. Histological examination identified only 2/152 (1.3%) endometrial tuberculosis while culture-proven endometrial TB was 2.6% (4/152) (14).

At our hospital, Saint Paul’s Hospital Millennium Medical College, endometrial biopsy was done twice per week as an outpatient office procedure with an average of 5-6 samples were taken per day. The aim of this study is therefore to determine the endometrial biopsy histopathological findings and associated factors at Saint Paul’s Hospital Millennium Medical College. The finding of the study could inform common endometrial pathology in developing country and associated clinical factors to tailor the management of patients based on documented evidences.

## Methods

### Study area and duration

This is a hospital-based retrospective study conducted at Saint Paul’s Hospital Millennium Medical College(SPHMMC) in Addis Ababa, the capital city of Ethiopia. Saint Paul’s hospital is one of the tertiary referral hospitals directly under the federal ministry of health. It is also a teaching hospital for the Millennium Medical College. The hospital offers service to 200,000 people annually who were referred from all corners of the country. On average 650 patients visit the hospital as outpatient and emergency daily. It gives service under different clinical disciplines which include obstetrics and gynecology (Statistics office of the hospital).

The study was a retrospective analysis of a cases of patients for whom endometrial biopsy was taken in SPHMMC, from January 2017 to January 2019. Patients for whom endometrial biopsy was taken during the study period and histopathologic result attached to patient’s file were included.

### Sample Size, sampling technique and data collection procedure

Sample size was determined using a single population proportion formula by taking P=0.236, the prevalence of secretory endometrium, the most common histopathologic finding in the previous study (13).

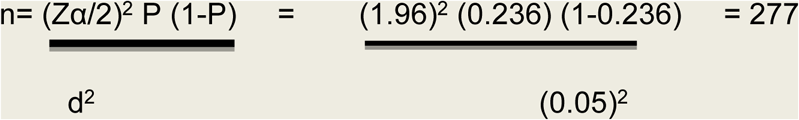

Where:

▪ n=the required sample size
▪ p= the proportion of the secretory endometrium in the previous study. (5).
▪ Zα/2=the critical value at 95% confidence level of certainty (1.96)
▪ d=the margin of error between the sample and the population 5%.

The registration numbers of cases for whom endometrial biopsy was done were identified from the procedure room register of the department of obstetrics and gynaecology. Their case files were retrieved from the medical record department of the hospital. A structured and pretested data collection proforma were used to abstract sociodemographic characteristics, gynecologic variables and histopathologic findings. Data abstraction was done by second-year residents after orientation by the principal investigator.

### Data processing and analysis

Data were entered to epi info version 3, then imported to and analyzed using SPSS version 21.0. Bivariate and multiple logistic regression analysis was done to see the association of variables. Descriptive statistics were used to characterize the dependent and independent variables. Chi-square test was used for statistical analysis of results. Odd’s ratios, 95% confidence interval and p-value set at 0.05 were used to determine the statistical significance of the associations.

### Independent variables

We considered independent variables like age, place of residence, marital status, parity contraceptive use, indications for biopsy and types of AUB.

### Dependent variables

We considered the following histopathologic findings: as dependent variables: Secretory endometrium, proliferative endometrium, atrophic endometrium, endometritis, endometrial polyp, endometrial hyperplasia,endometrial malignancy, complications of pregnancy, Inadequate sample and inconclusive result.

### Operational definitions

Normal histopathologic findings (normal cycling endometrium): are findings consistent with normal physiological menstrual patterns, i.e. proliferative and secretory endometrium.

Abnormal histopathologic findings: are findings other than normal physiologic menstrual changes. It includes hyperplasia, polyp, pregnancy complications, malignancy etc.

Pregnancy complications: are histopathologic findings of either molar or retained products of conception.

Inadequate sample: lack of any intact endometrial tissue specimen of adequate volume for the pathologist to identify endometrial pathology.

Inconclusive: histologic report by pathologists in cases where the finding doesn’t have specific pathologic features.

### Ethical consideration

Ethical clearance & permission letter to conduct the study and publish the outcome was obtained from the Institutional Review Board (IRB) of SPHMMC. Confidentiality was maintained during data collection, analysis and interpretation by avoiding recording of names and returning client records to its place after completion of data collection. All the datasets used and/or analyzed during the current study are included in the manuscript and available from the corresponding author on reasonable request.

## Result

### Socio-demographic and reproductive characteristics of study participants

A total of 361 lists of patient’s card number were retrieved from procedure room registry, out of which 301 patient records were obtained from the card room. Twenty-four of the records had incomplete data and they were excluded making the card retrieval rate of 83.3 %. A total of 277 patient records with complete data were analyzed. Majority of the women were in the age group of 40-49 years (32.13%) followed by 30-39 years (29.96) and ≥50 years (25.27%) with the mean age being 41.89 (Age range 20-85). One hundred eighty-nine (68.23%) of the patients were married and one hundred and fifty-seven (56.68%) of them were from the urban area. About 77.9% of the study participants were multi-parous and only 19.49 % were nulliparous. Forty-four (15.88%) of the participants had a history of at least one form of contraceptives use, the commonest type being used pills (65.91%).

Among all patients for whom the endometrial biopsy sample was taken during the study period, TVUS (transvaginal ultrasound) was done only for 6 (2.26%) of cases for endometrial assessment before the procedure. Majority of the samples were taken by an Intern (72.92%) and the commonest method of sampling was manual vacuum aspiration (MVA) (99.62%). Repeat sampling was done for 24 cases all due to inadequate prior (Table 1).

**Table 1:**
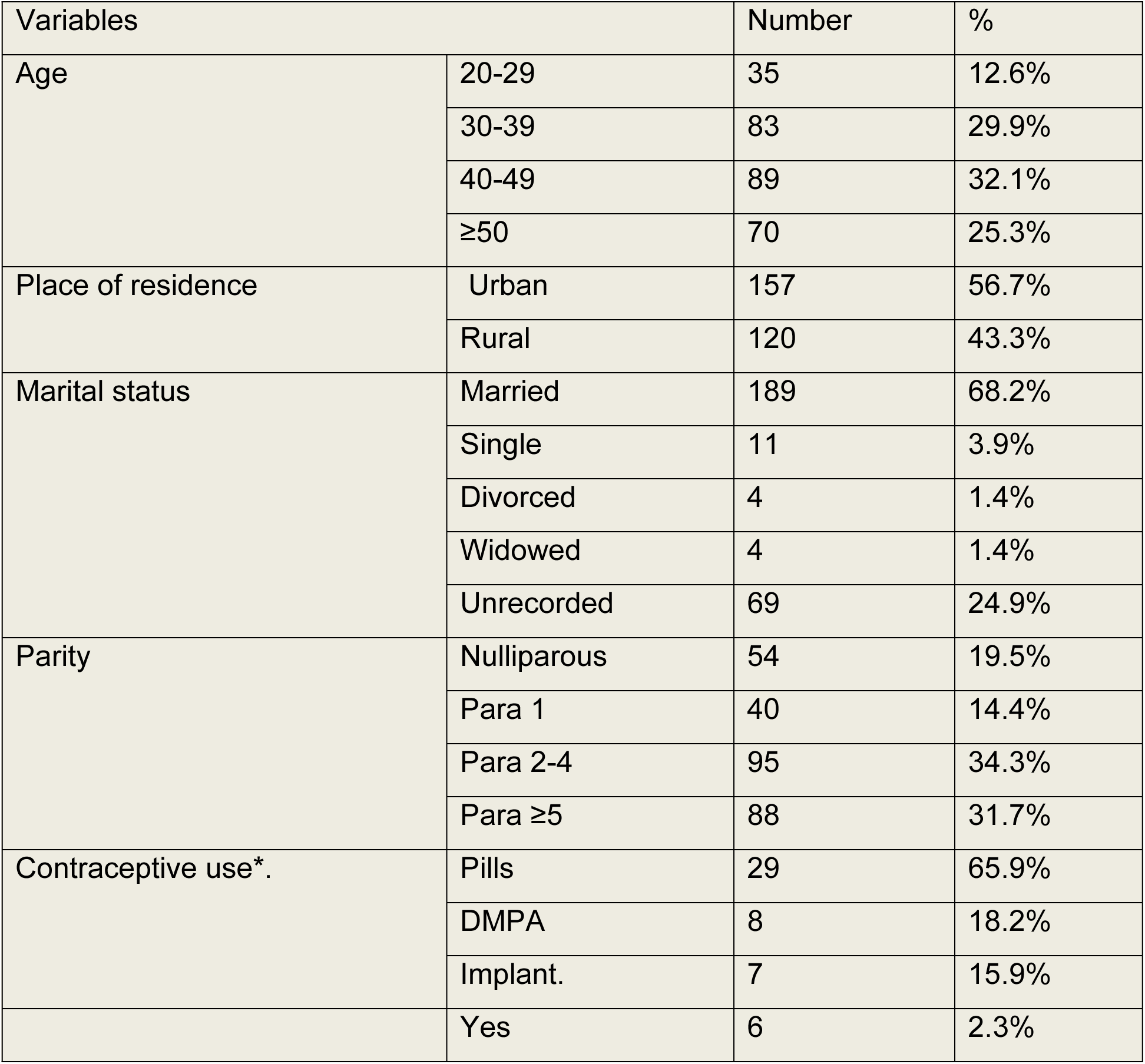
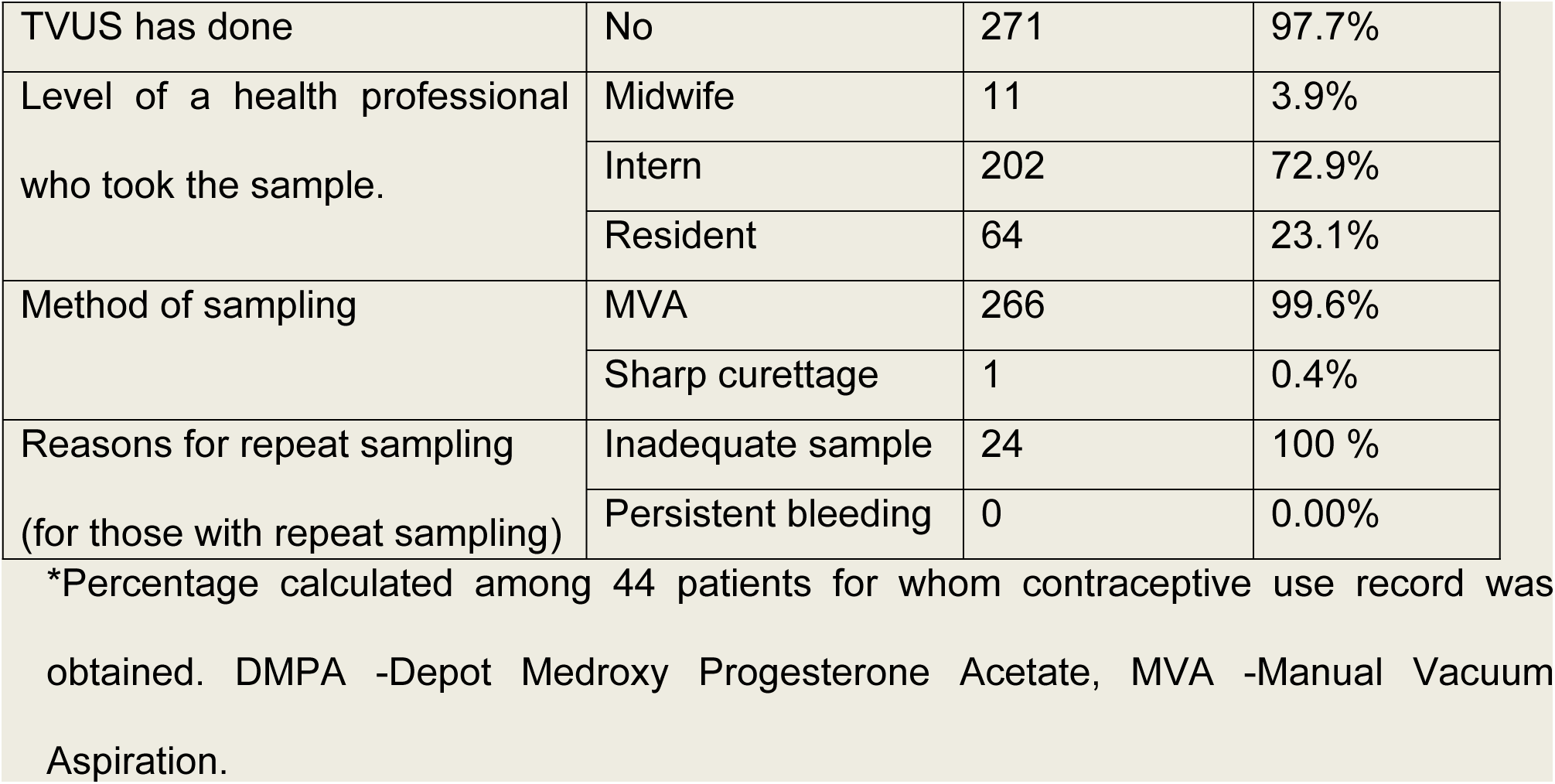
Socio-demographic and reproductive characteristics of patients for whom endometrial biopsy sample was taken at SPHMMC from February 2017 to January 2018, Addis Ababa, Ethiopia.

| Variables |  | Number | % |
| --- | --- | --- | --- |
| Age | 20-29 | 35 | 12.6% |
|  | 30-39 | 83 | 29.9% |
|  | 40-49 | 89 | 32.1% |
|  | ≥50 | 70 | 25.3% |
| Place of residence | Urban | 157 | 56.7% |
|  | Rural | 120 | 43.3% |
| Marital status | Married | 189 | 68.2% |
|  | Single | 11 | 3.9% |
|  | Divorced | 4 | 1.4% |
|  | Widowed | 4 | 1.4% |
|  | Unrecorded | 69 | 24.9% |
| Parity | Nulliparous | 54 | 19.5% |
|  | Para 1 | 40 | 14.4% |
|  | Para 2-4 | 95 | 34.3% |
|  | Para ≥5 | 88 | 31.7% |
| Contraceptive use*. | Pills | 29 | 65.9% |
|  | DMPA | 8 | 18.2% |
|  | Implant. | 7 | 15.9% |
|  | Yes | 6 | 2.3% |
| TVUS has done | No | 271 | 97.7% |
| Level of a health professional who took the sample. | Midwife | 11 | 3.9% |
|  | Intern | 202 | 72.9% |
|  | Resident | 64 | 23.1% |
| Method of sampling | MVA | 266 | 99.6% |
|  | Sharp curettage | 1 | 0.4% |
| Reasons for repeat sampling<br>(for those with repeat sampling) | Inadequate sample | 24 | 100 % |
|  | Persistent bleeding | 0 | 0.00% |

\*Percentage calculated among 44 patients for whom contraceptive use record was obtained. DMPA -Depot Medroxy Progesterone Acetate, MVA -Manual Vacuum Aspiration.

### Indication for Endometrial Biopsy

The commonest indication for endometrial biopsy was AUB in 239(86.28%) cases followed by thickened endometrium in 20(9.03%) and Infertility in 5 (1.81). The remaining 13 (4.69%) were done for pregnancy-related complications. Among all patients for whom endometrial biopsy sample was taken for an indication of AUB, the commonest AUB type was Intermenstrual bleeding (43.51%) followed by postmenopausal bleeding (27.19%) and heavy menstrual bleeding (23.01%) (Table 2).

**Table 2:** Frequency and percentage distribution of indications for Endometrial Biopsy of patients for whom endometrial biopsy sample was taken at SPHMMC from February 2017 to January 2018, Addis Ababa, Ethiopia.

|  |  |  |  |
| --- | --- | --- | --- |
| Indication for endometrial biopsy | AUB | 239 | 86.28% |
|  | -Heavy menstrual bleeding | 55 | 23.01% |
|  | -Intermenstrual bleeding | 104 | 43.51% |
|  | -Postmenopausal Bleeding | 65 | 27.19% |
|  | -Frequent menstrual bleeding. | 12 | 5.02% |
|  | -Others | 3 | 1.25% |
|  | Thickened endometrium | 20 | 7.22% |
|  | Infertility | 5 | 1.81% |
|  | Pregnancy related complications | 13 | 4.69% |
| Total |  | 277 | 100 |

### Histopathology Findings and associated factors

Out of total 277 cases of endometrial biopsy samples, the commonest histopathologic finding was endometrial polyp in 66 (23.83%) of cases followed by proliferative endometrium 47 (16.97%) and secretory endometrium 25(9.03%). Endometrial hyperplasia, endometrial malignancy, and pregnancy complications were reported in 9 (3.25%), 13 (4.69%), and 25 (9.03%) of cases respectively. Endometritis was detected in 20(7.22%), while Tuberculous (TB) endometritis was reported in 3(1.08%) of cases only. Inconclusive and inadequate sample was reported reports in 30 (10.83%) and 34 (12.27%) of cases respectively (Table 3).

**Table 3:**
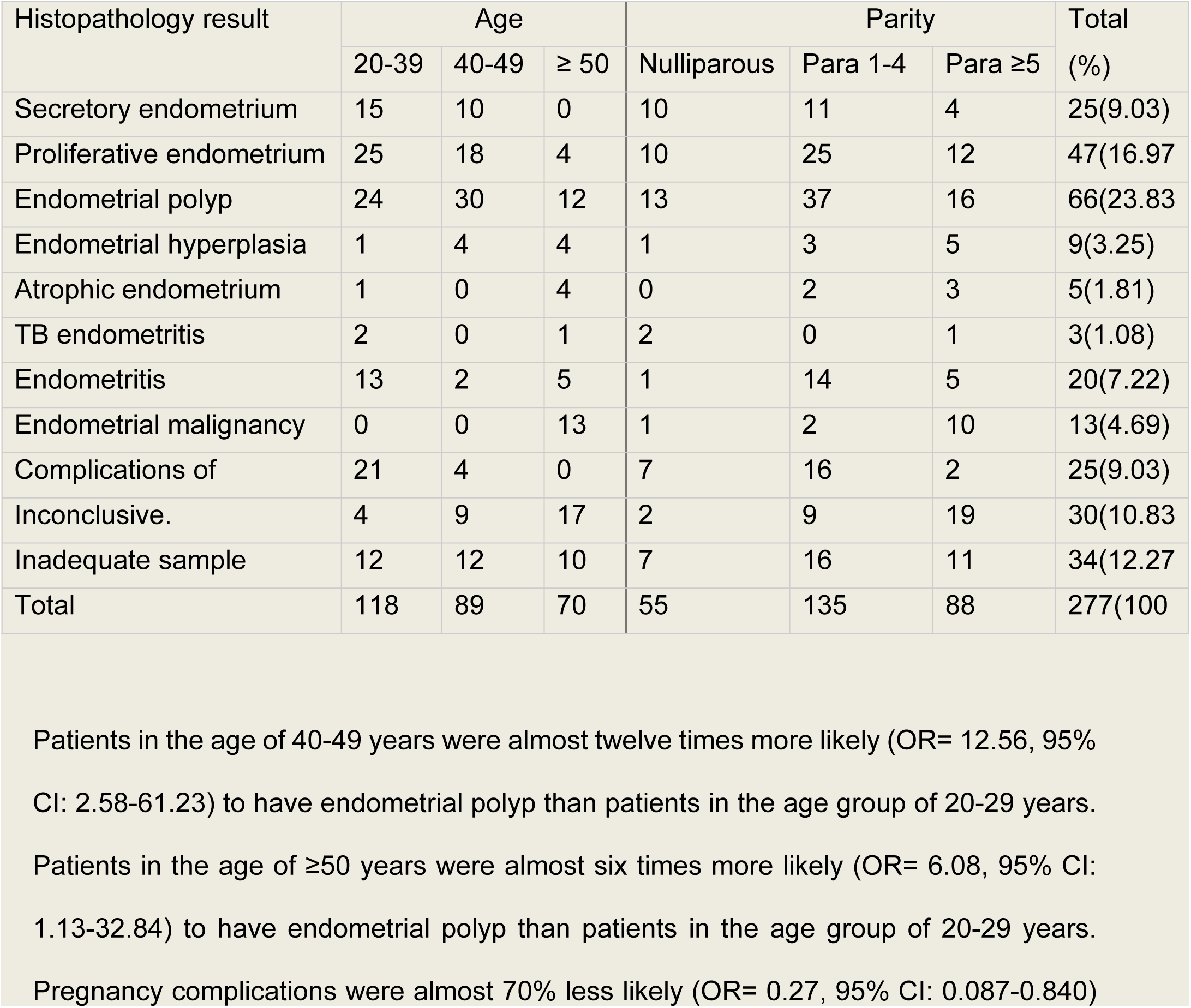
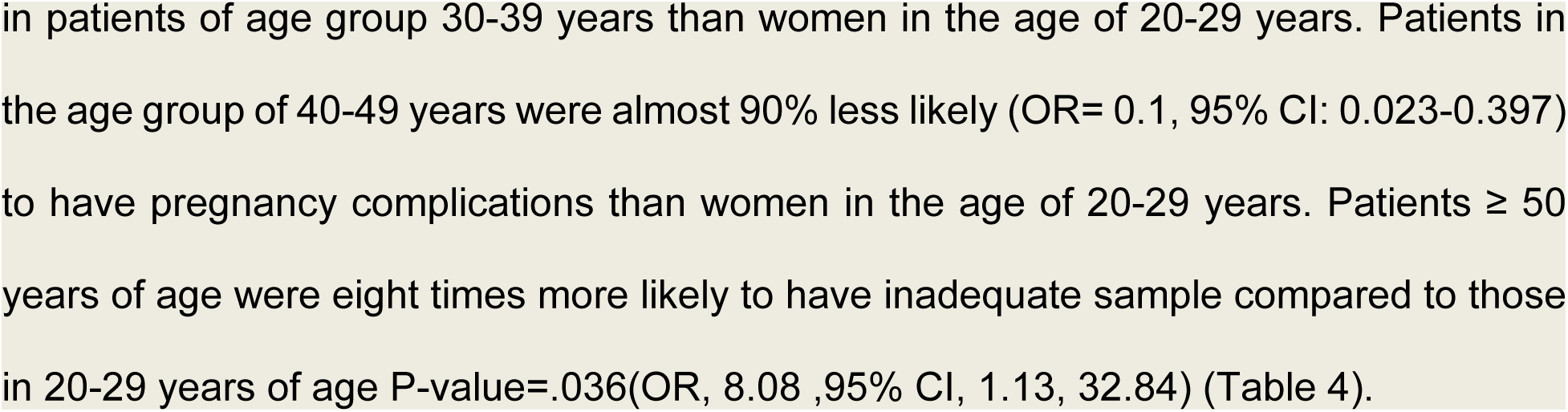
Frequency and percentage distributions of histopathology results for cases whom endometrial biopsy sample was taken at SPHMMC from February 2017 to January 2018, Addis Ababa, Ethiopia.

Eighty eight percent (88.8%) of endometrial hyperplasia were found in patients ≥40 years and 80% of atrophic endometrium in ≥50 years. Total of three cases with TB endometritis was found, among which two (66.6%) were detected in both nulliparous and each found in the age group of 20-29 years. All the cases with endometrial malignancy were found in the age of ≥50 years (Table 3).

Patients in the age of 40-49 years were almost twelve times more likely (OR= 12.56, 95% CI: 2.58-61.23) to have endometrial polyp than patients in the age group of 20-29 years. Patients in the age of ≥50 years were almost six times more likely (OR= 6.08, 95% CI: 1.13-32.84) to have endometrial polyp than patients in the age group of 20-29 years. Pregnancy complications were almost 70% less likely (OR= 0.27, 95% CI: 0.087-0.840) in patients of age group 30-39 years than women in the age of 20-29 years. Patients in the age group of 40-49 years were almost 90% less likely (OR= 0.1, 95% CI: 0.023-0.397) to have pregnancy complications than women in the age of 20-29 years. Patients ≥ 50 years of age were eight times more likely to have inadequate sample compared to those in 20-29 years of age P-value=.036(OR, 8.08, 95% CI, 1.13, 32.84) (Table 4).

**Table 4:** Association of Endometrial polyp, pregnancy complications and inadequate sample with selected characteristics of patients for whom endometrial biopsy sample was taken at SPHMMC, from February 2017 to January 2018, Addis Ababa, Ethiopia.

| Patient Characteristics |  | Endometrial polyp |  | P-Value | OR (95 % C. I) |
| --- | --- | --- | --- | --- | --- |
|  |  | No (%) | Yes (%) |  |  |
| Age in years | 20-29 (Ref) | 33 (94.3) | 2 (5.7) | .007 | - |
|  | 30-39 | 61 (73.5) | 22 (26.5) | .014 | 7.07 (1.48, 33.82) |
|  | 40-49 | 59 (66.3) | 30 (33.7) | .002 | 12.56 (2.58,61.23) |
| | $\geq 50$ | 58 (82.9) | 12 (17.1) | .036 | 6.08 (1.13, 32.84) |
| Parity | Nulliparous | 39 (75.0) | 15 (25.0) | .298 |  |
|  | Para 1 | 29 (76.3) | 10 (23.7) | .327 | 0.59 (0.21, 1.69) |
|  | Para 2-4 | 68 (73.1) | 25 (26.9) | .279 | 0.62 (0.26, 1.48) |
| | Para $\geq 5$ | 69 (81.2) | 16 (18.8) | .057 | 0.39 (0.15, 1.03) |
|  |  | Complications of pregnancy |  |  |  |
|  |  | No | Yes |  |  |
| Age in years | 20-29 (Ref) | 24 (68.6) | 11 (31.4) | .013 | - |
|  | 30-39 | 73 (88) | 10 (12) | .024 | 0.27 (0.087, 0.840) |
|  | 40-49 | 85 (95.5) | 4(4.5) | .001 | 0.1 (0.023, 0.397) |
| | $\geq 50$ | 70 (100) | 0 (0) | .997 | 0.000 |
| Parity | Nulliparous | 45 (86.5) | 7 (13.5) | .545 | - |
|  | Para 1 | 33 (86.8) | 5 (13.2) | .353 | 1.9 (0.49, 7.36) |
|  | Para 2-4 | 83 (89.2) | 10 (10.8) | .211 | 2.19 (0.641, 7.51) |
|  | Para ≥5 | 83 (97.6) | 2 (2.4) | .927 | 1.09 (0.17, 6.94) |
| Patient Characteristics |  | Inadequate sample. |  | P-Value | OR (95 % C. I) |
|  |  | No | Yes |  |  |
| Age in years | 20-29 (Ref) | 32(86.5) | 5(13.5) | .125 | - |
|  | 30-39 | 76(91.6) | 7(8.4) | .094 | 7.07 (1.48, 33.82) |
|  | 40-49 | 78(86.6) | 12(13.4) | .068 | 12.56 (2.58,61.23) |
|  | ≥ 50 | 56(84.8) | 10(15.2) | .036 | 8.08 (1.13, 32.84) |
| Parity | Nulliparous | 48(88.9) | 6(11.1) | .298 | - |
|  | Para 1 | 34(85) | 6(15) | .327 | 1.02 (0.21, 1.69) |
|  | Para 2-4 | 85(89.5) | 10(10.5) | .279 | 1.04 (0.26, 1.48) |
|  | Para ≥5 | 76(86.4) | 12(13.6) | .067 | 0.39 (0.15, 1.03) |
| Health professional<br>who took sample. | Midwife | 9(81.8) | 2(18.2) | .065 | - |
|  | Intern | 180(89.1) | 22(10.9) | .172 | 0.62(0.50-4.92) |
|  | Resident | 54(84.4) | 10(15.6) | .357 | 0.45(0.26-5.85) |

## Discussion

In the current study, the mean age of the study patients was 41.89 years with a minimum value of 20 and a maximum of 85 years. It shows older age group compared to study at Tikur Anbessa Specialized Teaching Hospital(TASTH), which showed the mean age of 36.75 years (13).

In our study, the commonest indication for endometrial biopsy was AUB in 239(86.28%) cases followed by thickened endometrium, 20(7.22%) and pregnancy-related complications,13 (4.69%). Indications were comparable to the study at TASTH which showed AUB in 60.9%, followed with Infertility, lower abdominal/pelvic pain and pregnancy-related complications accounting for 10.9%,10.7% and 9.4% respectively (13).

Endometrial polyp was the commonest histologic finding in current study accounting for 23.8% of cases. This finding is comparable with study in Romania in which endometrial polyp accounts for 22.2% of the endometrial pathology (12). The prevalence of endometrial polyp was higher in the current study compared to studies in Sudan and Ethiopia which account for 15.2% in both studies(13, 15). This high prevalence can be explained with the high number of patients in the age group 40-49 years (32.13%) in our study compared to study at TASTH which was 25.7%(13). Patients in the age of 40-49 years were 12 times more likely (OR= 12.56, 95% CI: 2.58-61.23) to have endometrial polyp. The finding of proliferative endometrium (17.0%) was higher compared to study in Sudan (9.5%) and it was comparable to study at TASTH (18.1%)(13, 15).

The prevalence of pregnancy complications (9.03%) in the current study was lower compared to 17.9% and 62.6% reports from the study in Sudan and Nigeria respectively (15, 16) but was comparable to 10.8% report of TASTH study finding (13).

Endometrial hyperplasia was detected in 9 (3.25%) cases while endometrial malignancy in 13 (4.69%) of cases in the current study. The prevalence of both was much lower compared to the study done in Romania in which Hyperplasia’s and Endometrial malignancy were reported in 54.9% and 19% of the patients respectively (12). The finding of Endometrial hyperplasia was comparable to the findings in Sudan and Ethiopia which shows a prevalence of 2.8% and 3.1% respectively. Endometrial malignancy was higher compared with study at TASTH (1%) and its lower compared with study in Sudan (8.9%) (13, 15). Higher rate of endometrial malignancy when compared to study at TASTH was explained by a higher rate of the older age group of ≥50 years in the current study,9.8% versus 25.27%(13).

In the current study TB endometritis was detected in 3(1.08%) which was lower than the prevalence from the study in India, showing cases of tuberculous endometritis in 3(3%) (17), but comparable to studies in Nigeria and Ethiopia in which histological examination identified 3(0.9%) and 2(1.3%) of endometrial tuberculosis respectively(14, 18).

In our study samples were reported as inconclusive and inadequate in 30 (10.83%) and 34 (12.27%) of cases respectively. The prevalence of inadequate sample was high compared to a meta-analysis of 39 studies involving 7914 women, showing only fewer than 5 percent of patients had an insufficient or no sample(7). Inconclusive results were also higher compared to study at TASTH which accounts for 7.3% cases(13, 19). The inadequate sample was comparable to studies in Nepal (12%) and Sudan (11.2%) (13, 20). Combining both inadequate and inconclusive reports in our study sample inadequacy was higher in the current study compared to all other publications which might be because of less endometrial assessment by transvaginal ultrasound. Retrospective review at the Khon Kaen University’s Srinagarind Hospital showed menopausal status and endometrial thickness of less than 8 mm were independent predictors for insufficient endometrial tissue after endometrial sampling(21). In our study patients ≥ 50 years of age were eight times more likely to have inadequate sample compared to those 20-29 years of age but TVUS was not done to assess endometrium in almost all cases (97.74%) and there is no association with parity and level of health professional who did the procedure. Office endometrial sampling using a pipelle device is most reliable and acceptable by the patient(1, 3). A study in Thailand for comparison of MVA versus Pipelle device shows better tissue adequacy in pipelle group (85.6% versus 81.1%). The study also highlighted higher difficult procedure with MVA (27.0% versus 14.4%) and significantly higher moderate to severe pain with MVA (60.7% versus 19.5%) (6). In the current study all the samples including the 24 repeat samples were taken using MVA which might have exaggerated inadequate sample.

## Conclusion

The commonest indication for endometrial biopsy was AUB, followed by thickened endometrium and pregnancy-related complications. Endometrial polyp was the commonest histopathologic finding followed by proliferative and secretory endometrium respectively. Patients in the age of 40-49 years were 12 times more likely to have endometrial polyp. Rate of sample inadequacy is higher than most of the study reports and patients ≥ 50 years of age were eight times more likely to have inadequate sample This mandates to improve sampling technique to increase sample adequacy by using more preferred methods like Pipelle device as an alternative to MVA and considering options of D & C and hysteroscopy especially in cases difficulty to get sample and in those with the prior inadequate sample. Routine transvaginal ultrasound for endometrial thickness evaluation especially for postmenopausal women is very important to decide the necessity of endometrial sampling. The current study has limitation as its retrospective document review and difficult to get some clinical history to determine factors associated with histopathologic findings. So, we recommend well designed prospective study to generate more strong evidence that guides patient management.

## List of abbreviations

ACOG: American College of Obstetricians and Gynecologists
AUB: Abnormal uterine bleeding
COC: Combined Oral Contraceptive
D&C: Dilation and curettage
DMPA: Depot medroxyprogesterone acetate
EMB: Endometrial biopsy
IUCD: Intrauterine contraceptive device
MVA: Manual Vacuum Aspiration
OCP: Oral Contraceptive Pills
RGOPD: Regular Gynecology Outpatient Department
SPHMMC: St. Paul’s Hospital Millennium Medical College
SPSS: Statistical package for social sciences
TASTH: Tikur Anbessa Specialized Teaching Hospital
TVUS: Transvaginal Ultrasonography

## Authors contributions

SK conceived the study, supervised data collection and drafted the manuscript. LBT, SK and MN performed the statistical analysis.ES supervised the whole process of the study from conception to preparation of draft of manuscript. LBT and GTF revised and edited the manuscript for publication. All authors revised and approved the final manuscript.

## Funding

This project did not get any funding support from any organization.

## Acknowledgement

We thank midwives at procedure room and workers of card room for providing us with procedure registration and patient record facilitating data collection.

## Competing Interests

The authors declare that they have no competing interests.

## References

1. ACOG. Practice Bulletin No. 149: Endometrial cancer. Obstetrics and gynecology. 2015;125(4):1006–26.

2. World Health Organization, Reproductive Health Research, Medical management of abortion %U http://www.ncbi.nlm.nih.gov/books/NBK5367792018.

3. Hoffman BL, Schorge JO, Bradshaw KD, Halvorson LM, Schaffer JI, Corton MM. Abnormal Uterine Bleeding. Williams Gynecology. 3 ed. New York, NY: McGraw-Hill Education; 2016.

4. Saadia A, Mubarik A, Zubair A, Jamal S, Zafar A. Diagnostic accuracy of endometrial curettage in endometrial pathology. Journal of Ayub Medical College Abbottabad. 2011;23(1):129–31.

5. Telner DE, Jakubovicz D. Approach to diagnosis and management of abnormal uterine bleeding. Canadian Family Physician. 2007;53(1):58–64.

6. Wanijasombutti P, Imruetaicharoenchok A, Tangjitgamol S, Loharamtaweethong K, Phuriputt N, Phaloprakarn C. Comparison of tissue adequacy for histologic examination from Ipas MVA plus and Wallach Endocell in women with abnormal uterine bleeding. Journal of Obstetrics and Gynaecology Research. 2015;41(8):1246–54.

7. Visser NCM, Reijnen C, Massuger L, Nagtegaal ID, Bulten J, Pijnenborg JMA. Accuracy of Endometrial Sampling in Endometrial Carcinoma: A Systematic Review and Meta-analysis. Obstetrics and gynecology. 2017;130(4):803–13.

8. Kisielewski F, Gajewska ME, Marczewska MJ, Panek G, Wielgos M, Kaminski P. Comparison of endometrial biopsy and postoperative hysterectomy specimen findings in patients with atypical endometrial hyperplasia and endometrial cancer. Ginekologia polska. 2016;87(7):488–92.

9. Shukla M, Fonseca MN, Kharat D, Tekale P. A study to correlate histopathological findings in patients with abnormal uterine bleeding. 2017. 2017;6(2):4.

10. Sirimai K, Lertbunnaphong T, Malakorn K, Warnnissorn M. Comparison of Endometrial Pathology between Tissues Obtained from Manual Vacuum Aspiration and Sharp Metal Curettage in Women with Abnormal Uterine Bleeding. 2016;99(2):8.

11. Ülgüt C, Paiusan L, Furau G, Stănescu C. Endometrial histophatology findings in postmenopausal women. Jurnal Medical Aradean (Arad Medical Journal). 2015;18(1):56–60.

12. Bohiltea R, Sajin M, Furtunescu FL, Bohiltea L, Mihart AE, Baro? A, et al., editors. Clinical and pathological correlations in endometrial pathology2015 2015.

13. Tukue D, Gebremariam L. Patterns of endometrial pathology at Tikur anbessa specialized teaching hospital, Addis Ababa university, Ethiopia. A retrospective study.

14. Abdissa S, Abebe T, Ameni G, Teklu S, Bekuretsion Y, Abebe M, et al. Endometrial tuberculosis among patients undergoing endometrial biopsy at Tikur Anbesa specialized hospital, Addis Ababa, Ethiopia. BMC Infectious Diseases. 2018;18(1):304.

15. Ali MH, Khalid IO, Khier MM, Adam NM. Histopathological Pattern of Endometrial Sampling in Abnormal Uterine Bleeding.

16. Asuzu IM, Olaofe OO. Histological Pattern of Endometrial Biopsies in Women with Abnormal Uterine Bleeding in a Hospital in North Central Nigeria. International Journal of Reproductive Medicine. 2018;2018:5.

17. Tiwari A, Kaur N, Jain S, Rai R, Jain SK. Histopathological Study of Endometrial Biopsy Specimens for Abnormal Uterine Bleeding. Journal of Lumbini Medical College. 2016;4(2):72–6.

18. Dike. E, Ojiyi. E, Department of Obstetrics and Gynaecology ASUtHAS-EN. Histopathological characteristics ofendometrial biopsies in the Igbos of Nigeria. J Med Res. 2017;3(4):187–90.

19. Hanegem Nv, Prins MMC, Bongers MY, Opmeer BC, Sahota DS, Mol BWJ, et al. The accuracy of endometrial sampling in women with postmenopausal bleeding: a systematic review and meta-analysis. European Journal of Obstetrics and Gynecology and Reproductive Biology. 2016;197:147–55.

20. Sharma S, Makaju R, Shrestha S, Shrestha A. Histopathological Findings of Endometrial Samples and its Correlation Between the Premenopausal and Postmenopausal Women in Abnormal Uterine Bleeding. Kathmandu University medical journal (KUMJ). 2014;12(48):275–8.

21. Aue-Aungkul A, Kleebkaow P, Kietpeerakool C. Incidence and risk factors for insufficient endometrial tissue from endometrial sampling. Int J Womens Health. 2018;10:453–7.

